# Gelation of Uniform Interfacial Diffusant in Embedded 3D Printing

**DOI:** 10.1101/2023.04.02.535250

**Authors:** Sungchul Shin, Lucia G. Brunel, Betty Cai, David Kilian, Julien G. Roth, Alexis J. Seymour, Sarah C. Heilshorn

## Abstract

While the human body has many different examples of perfusable structures with complex geometries, biofabrication methods to replicate this complexity are still lacking. Specifically, the fabrication of self-supporting, branched networks with multiple channel diameters is particularly challenging. Here, we present the Gelation of Uniform Interfacial Diffusant in Embedded 3D Printing (GUIDE-3DP) approach for constructing perfusable networks of interconnected channels with precise control over branching geometries and vessel sizes. To achieve user-specified channel dimensions, this technique leverages the predictable diffusion of crosslinking reaction-initiators released from sacrificial inks printed within a hydrogel precursor. We demonstrate the versatility of GUIDE-3DP to be adapted for use with diverse physiochemical crosslinking mechanisms by designing seven printable material systems. Importantly, GUIDE-3DP allows for the independent tunability of both the inner and outer diameters of the printed channels and the ability to fabricate seamless junctions at branch points. This 3D bioprinting platform is uniquely suited for fabricating lumenized structures with complex shapes characteristic of multiple hollow vessels throughout the body. As an exemplary application, we demonstrate the fabrication of vasculature-like networks lined with endothelial cells. GUIDE-3DP represents an important advance toward the fabrication of self-supporting, physiologically relevant networks with intricate and perfusable geometries.

## 1. INTRODUCTION

Channels with intricate geometries are critical for the function of many tissues throughout the human body, including vascular networks, lymphatic vessels, airway channels, and the gastrointestinal tract.^[1–5]^ These naturally evolved, perfusable geometries have inspired bioengineers to fabricate mimetic structures for use in organ-on-a-chip models^[6–8]^, as tissue engineered therapies^[9,10]^, and for studying the role of physical forces in regulating healthy and diseased cellular function^[11,12]^.

Compared to micromolding or microfluidics, 3D printing of biomaterial inks has emerged as a technique to create perfusable networks with user-specified channel paths that can be easily reconfigured.^[13,14]^ In one bioprinting strategy for channel formation, a tube-like structure is created through direct layer-by-layer printing of the channel walls.^[15–17]^ While this allows for fabrication of branched networks, the structures are prone to leakage due to insufficient adhesion between layers, and the resulting surfaces and interfaces are typically not smooth. To address these limitations, coaxial extrusion can be used to fabricate perfusable channels by directly printing hollow filaments from concentric nozzles.^[18–20]^ However, since the inner and outer diameters are set by the nozzle geometry, the final printed structure cannot include interconnected branch points. This technique is therefore limited to single-lumen geometries of fixed diameter. As an alternative approach, extrusion printing of a sacrificial ink enables patterning of void spaces within a bulk material to form interconnected lumens with smooth inner surfaces.^[21–23]^ In contrast to core-shell extrusion, this sacrificial ink strategy results in perfusable structures without a vessel-like shell. As such, this method cannot be used to form self-supporting networks.

To enable the biofabrication of perfusable, self-supporting channels with interconnected branch points and predictable, user-specified diameters, we developed a new strategy we term Gelation of Uniform Interfacial Diffusant in Embedded 3D Printing (GUIDE-3DP). We demonstrate the versatility of this platform to be adapted for multiple materials and physiochemical hydrogel crosslinking mechanisms, the independent tunability of both the inner tube diameter and outer shell diameter of printed channels, and the ability to fabricate seamless junctions at branch points. Finally, to exemplify the use of GUIDE-3DP for creating physiologically relevant structures, networks of perfusable channels are fabricated with an endothelial cell lining to mimic vasculature.

## 2. RESULTS AND DISCUSSION

Embedded 3D printing is an additive manufacturing strategy in which inks are extruded within a support material, decreasing deformation due to gravity or surface tension and thus enabling the printing of inks into complex geometries.^[24,25]^ In our GUIDE-3DP technique, we leverage the predictable diffusion of crosslinking reaction-initiators, released from sacrificial inks within the support material, to rapidly create perfusable, self-supporting networks with precise control over the branching geometry and vessel dimensions (**Figure 1**). In this strategy, a crosslinkable gel precursor is employed both as the support matrix for microextrusion printing as well as the material that eventually comprises the vessel walls. A sacrificial ink containing a cytocompatible reaction-initiator is printed into the gel precursor. Following printing, the reaction-initiators diffuse radially and uniformly across the ink interface and into the surrounding gel precursor (**Figure 1a, left panel**). The crosslinking reaction occurs only in regions where the gel precursor is in contact with the reaction-initiator, which can be selected to either induce spontaneous crosslinking or to require an external trigger such as light activation (**Figure 1a, middle panel**). Finally, the sacrificial ink and uncrosslinked support material are removed, resulting in a self-supporting, perfusable printed structure (**Figure 1a, right panel**).

**Figure 1.**
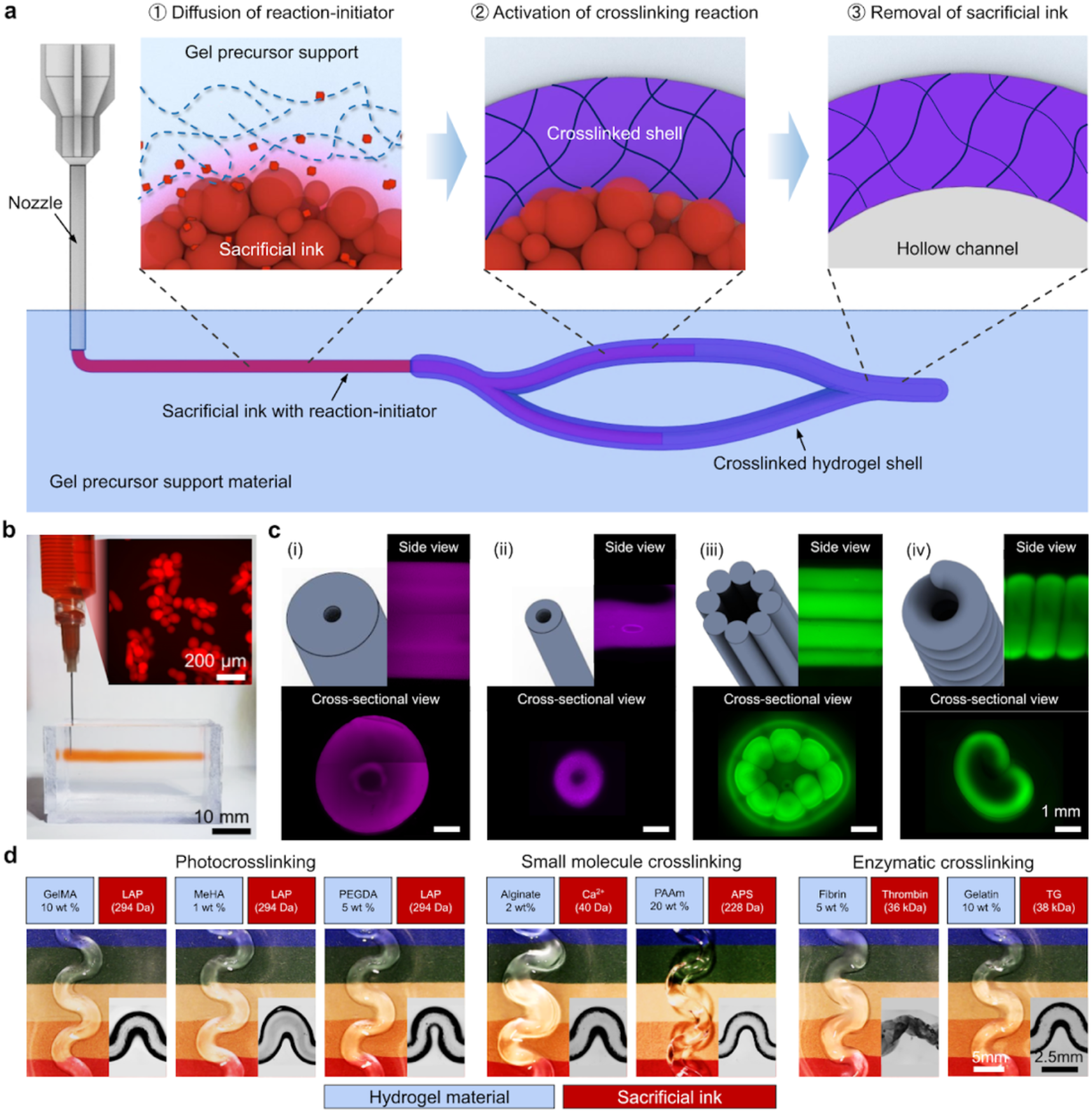
The Gelation of Uniform Interfacial Diffusant in Embedded 3D Printing (GUIDE-3DP) method. **(a)** Sequence of key steps in the GUIDE-3DP method: (1) A sacrificial ink containing a reaction-initiator is extruded into a corresponding gel precursor support material, and the reaction-initiator diffuses out of the sacrificial ink and into the gel precursor. (2) The crosslinking reaction occurs in areas that contain both the gel precursor and the reaction-initiator, forming a shell of crosslinked gel. (3) The sacrificial ink and unreacted gel precursor material are removed, resulting in a printed structure with an open, hollow lumen. **(b)** A sacrificial ink composed of gelatin microparticles (inset) is extruded into a GelMA support material. **(c)** The GUIDE-3DP method (panels i, ii) allows for the fabrication of seamless hollow channels, unlike the conventional approach of direct, layer-by-layer printing of tubular constructs (panels iii, iv). **(d)** The GUIDE-3DP method is amenable to a variety of common biomaterials and crosslinking approaches, demonstrated here with photocrosslinking (GelMA/LAP, MeHA/LAP, PEGDA/LAP), small molecule crosslinking (alginate/calcium ions (Ca^2+^), PAAm/ammonium persulfate (APS)), and enzymatic crosslinking (fibrin/thrombin, gelatin/transglutaminase (TG)).

As a first demonstration, light-crosslinkable gelatin methacryloyl (GelMA) was selected as the gel precursor support material with a slurry of gelatin microparticles loaded with a photoinitiator (lithium phenyl-2,4,6-trimethylbenzoylphosphinate, LAP) as the sacrificial ink. When a filament of the sacrificial ink is printed within the GelMA support material (**Figure 1b**), the encapsulated reaction-initiator (*i*.*e*., LAP) begins to diffuse radially away from the ink. Upon LAP activation with ultraviolet (UV) light, a continuous shell of crosslinked GelMA is formed around the sacrificial ink, which can then be removed after melting, resulting in a perfusable channel (**Figure 1c, panels i and ii**). In contrast to the GUIDE-3DP approach, previous layer-by-layer demonstrations of printing self-supporting channels typically result in surfaces that are discontinuous and prone to defects that may cause leakage due to gaps or delamination of the additive layers (**Figure 1c, panels iii and iv**). Moreover, these traditional additive manufacturing strategies are time-intensive, since multiple passes are required to print a single channel, and limited in their ability to tune the channel dimensions, since the wall thickness cannot be less than the thickness of a single printed filament. GUIDE-3DP overcomes all three of these constraints. The channel surface is uniform since the diffusion of the reaction-initiator into the support material is uniform; the total print time is reduced since the nozzle moves along the print path only once for a single channel; and the shell thickness is controlled by setting the diffusion time. Moreover, the GUIDE-3DP strategy is amenable to a broad range of polymers and crosslinking mechanisms. To demonstrate this, we formulated seven different gel precursor materials to enable crosslinking by the following mechanisms: (1) photocrosslinking with UV light for GelMA, hyaluronic acid methacrylate (MeHA), and poly(ethylene glycol) diacrylate (PEGDA), (2) crosslinking with small molecules for alginate and polyacrylamide (PAAm), and (3) enzymatic crosslinking for fibrin and gelatin. All seven distinct support material/sacrificial ink combinations were successfully used for GUIDE-3DP fabrication of perfusable structures (**Figure 1d**). A table detailing the material composition and printing parameters for all constructs in this manuscript is presented in **Table S1**.

The GUIDE-3DP strategy is enabled by the material properties of the sacrificial ink and gel precursor support material. As the hydrogel component of the sacrificial inks for GUIDE-3DP, we chose a slurry of gelatin microparticles (**Figure 2**). The gelatin microparticles (mean diameter of 18.0 ± 4.0 µm) were fabricated with complex coacervation and concentrated by filtration to form a jammed microparticle ink (**Figure 2a-b**). After comparing the rheological analysis of concentrations ranging from 4-10 wt%, the 8% gelatin microgel ink was identified as having the viscoelastic properties required for printing smooth filaments within the GelMA support material (**Figure S1**). Specifically, the 8 wt% gelatin microparticle slurry demonstrated important sacrificial ink characteristics, including (1) yield stress and shear-thinning behavior (**Figure 2c-d**), which enables extrusion through the print nozzle; (2) rapid self-healing (**Figure 2e**), which limits ink spreading and improves print resolution; and (3) controllable sol-gel phase transition, melting at temperatures above 34 °C (**Figure 2f**), which facilitates its removal from the final print.

**Figure 2.**
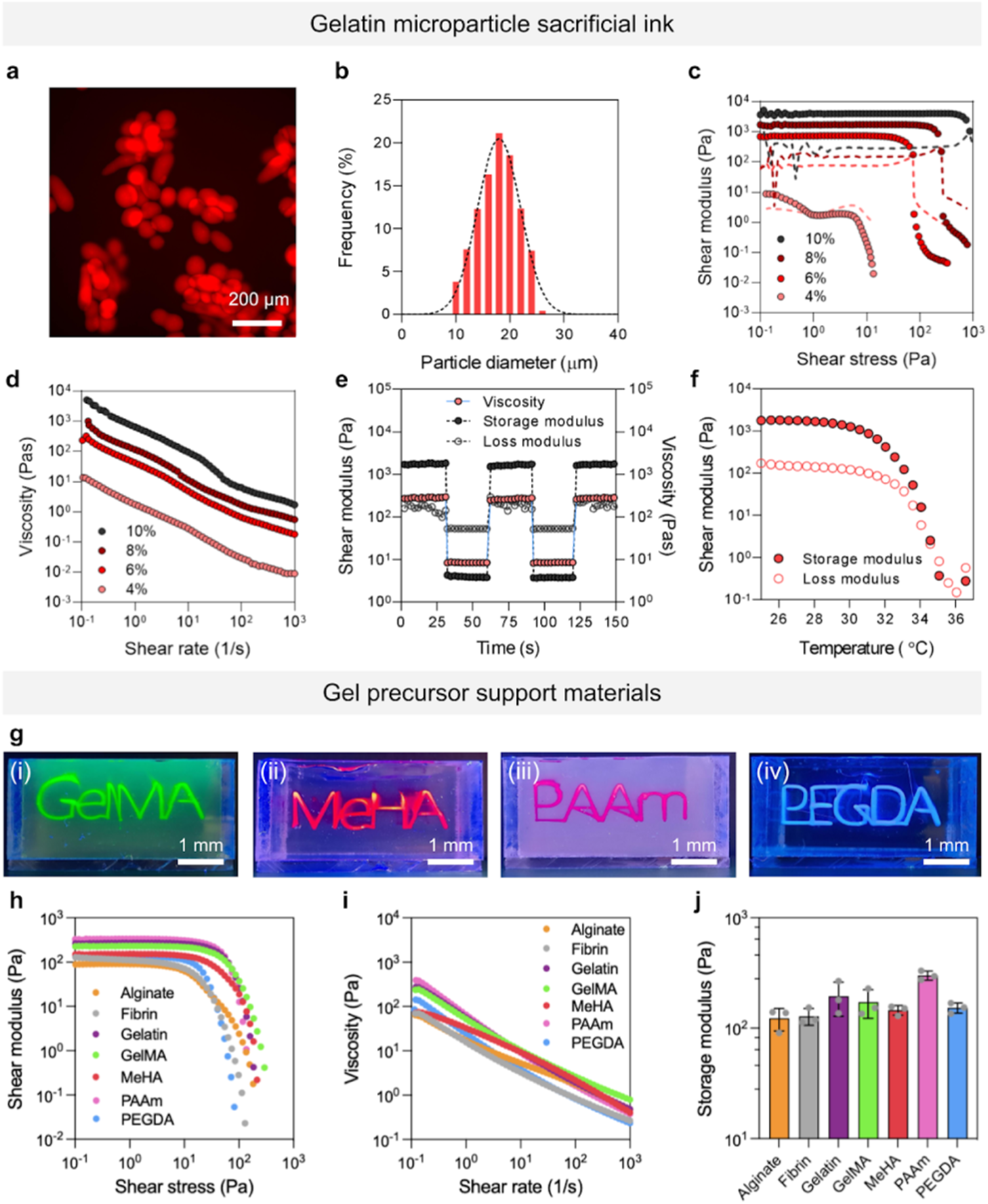
Material properties of the sacrificial ink and gel precursor support materials that enable GUIDE-3DP. **(a)** Fluorescence images of the gelatin microparticles used in the sacrificial ink prior to jamming, stained with rhodamine. **(b)** Distribution of gelatin microparticle diameters. Formulations of the gelatin microparticle sacrificial inks ranging from 4-10 wt% exhibit **(c)** yield stress (storage modulus G’ represented with filled circles; loss modulus G’’ represented with dashed lines) and **(d)** shear-thinning behavior. The gelatin microparticle sacrificial ink (8 wt%) exhibits **(e)** self-healing and **(f)** thermoreversible behavior. **(g)** Sample prints of selected gel precursor support materials (i) GelMA, (ii) MeHA, (iii) PAAm, and (iv) PEGDA. Each gel precursor support material (alginate, PAAm, GelMA, MeHA, PEGDA, fibrin, and gelatin) with 2 wt% of the viscosity modifier Aristoflex AVC added has **(h)** a yield stress, **(i)** shear-thinning behavior, and **(j)** storage moduli between 100-1000 Pa in the linear viscoelastic regime.

We next demonstrated that by changing the reaction-initiator within the sacrificial ink, we can construct prints with a variety of gel precursors and crosslinking mechanisms using GUIDE-3DP (**Figure 2g**). We achieved suitable rheological properties of each gel precursor support material (alginate, PAAm, GelMA, MeHA, PEGDA, fibrin, and gelatin) by adding Aristoflex Ammonium Acryloyldimethyltaurate/VC Copolymer (AVC) as a viscosity modifier (2 wt%) (**Figures 2h-j; S2**). For each crosslinking system, a bespoke set of reaction-initiator molecules was designed to diffuse across the interface between the printed ink and the support material (**Table S1**). Specifically, for the photocrosslinkable gel precursor support materials, the sacrificial gelatin microparticle ink was loaded with LAP (294 Da) as a photoinitiator, while Ca^2+^ (40 Da) was loaded as the small molecule for alginate, a cation-crosslinked gel precursor material. As PAAm requires both polymerization and crosslinking to occur simultaneously, the polymerization initiator ammonium persulfate (APS, 228 Da) and the crosslinker bis-acrylamide were included in the gelatin sacrificial ink and acrylamide support material, respectively. For the enzymatically crosslinked materials, the sacrificial ink was loaded with the enzyme thrombin (36 kDa) or transglutaminase (38 kDa) to induce gelation of fibrin or gelatin, respectively. Importantly, to avoid the transglutaminase-mediated crosslinking of the sacrificial ink for the gelatin support material, the sacrificial ink used was Pluronic F-127 (24 wt%) instead of gelatin microparticles. Pluronic F-127 is a thermoreversible triblock copolymer commonly used as a sacrificial hydrogel in 3D bioprinting applications.^[21,23,26–28]^

Both the inner and outer shell diameters of GUIDE-3DP channels are highly tunable, since the inner diameter is the same as the diameter of the printed sacrificial ink filament, and the outer shell diameter is dependent on the diffusion time (**Figure 3**). GUIDE-3DP offers two strategies for dynamically varying the ink filament diameter during direct writing. In one strategy, the applied dosing pressure is varied while the writing speed is held constant (**Figure 3a-b**); in the other strategy, the applied pressure is held constant while the writing speed is varied (**Figure 3c**). In both cases, the diameter of the printed ink filament dictates the inner diameter of the crosslinked GelMA tubes. The ability to control the inner diameter with the GUIDE-3DP strategy enables facile fabrication of biologically inspired, complex perfusable structures. For example, many perfusable structures in nature have inner diameters that are not constant, which can be replicated using the GUIDE-3DP strategy. As a demonstration, we created a model of the large intestine, which has a single lumen that is segmented into “pouch-like” haustra (**Figure 3d**), by alternating the print speed of the sacrificial ink (1 mm/s, 40 psi for an inner diameter of 4.5 mm; 2 mm/s, 40 psi for an inner diameter of 3.5 mm). As a second demonstration, we printed a perfusable network with both diverging and converging vessel branch points that emulate the natural branching observed in vascular networks *in vivo*, with parent vessels of larger inner diameter (2.5 mm) and daughter vessels of smaller inner diameter (1.5 mm) (**Figure 3e**). Upon gelatin melting and extraction of the fabricated GelMA structures, both the large intestine model and the vascular network model were readily perfusable with dye.

**Figure 3.**
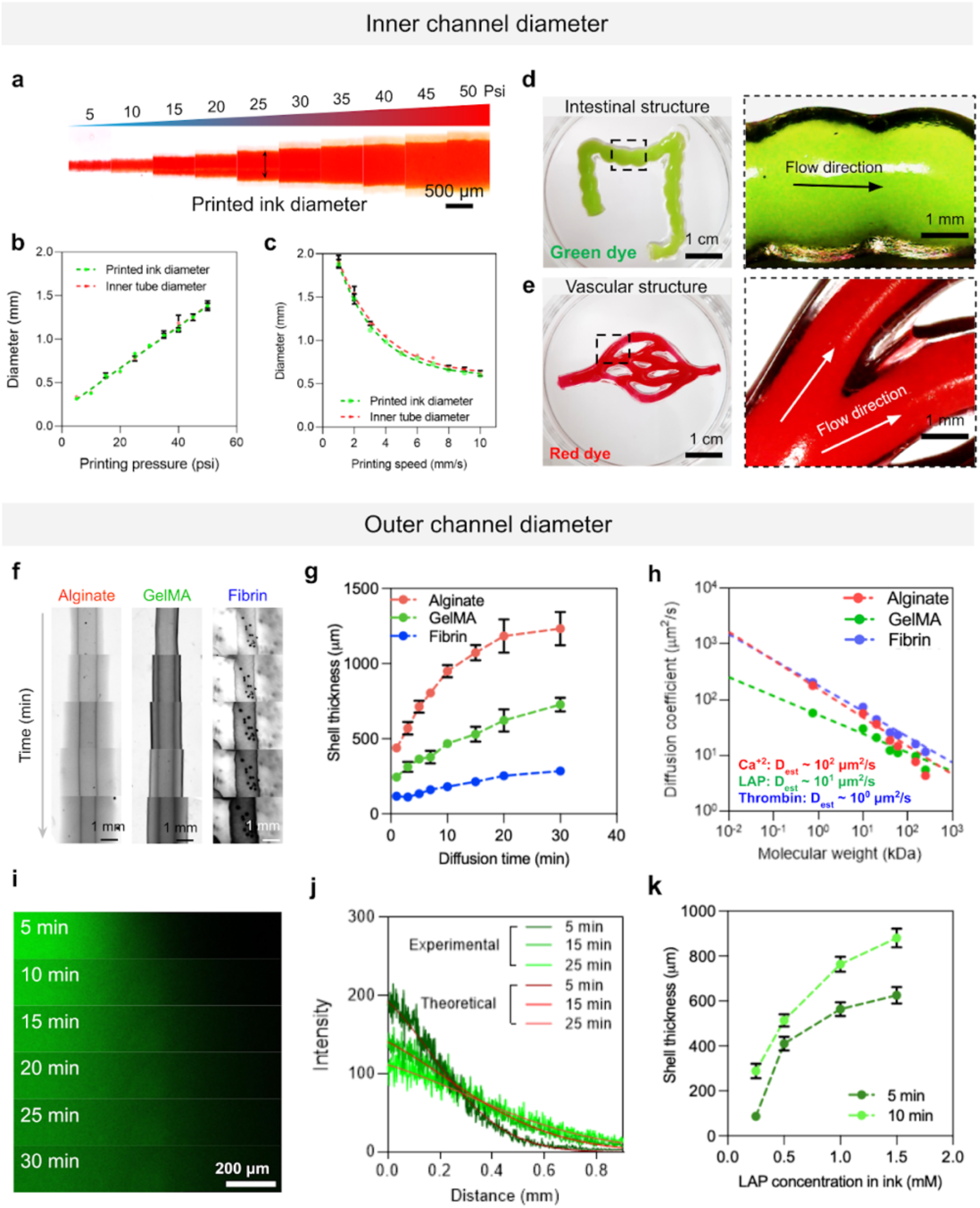
The inner and outer diameters of channels printed with GUIDE-3DP can be independently tuned. **(a)** The diameter of printed inks increases with the application of greater pressure during extrusion, demonstrated between 5-50 psi. The inner diameter of the crosslinked channel corresponds to the diameter of the printed ink filament, which can be controlled **(b)** as a function of applied pressure (demonstrated at a constant printing speed of 5 mm/s) and **(c)** as a function of printing speed (demonstrated at a constant applied pressure of 25 psi). **(d)** An intestinal model of the haustra structure was printed by adjusting the printing speed alternately between 1 and 2 mm/s. In this way, the inner diameter of the channel undulated between 4.5 mm and 3.5 mm. Green dye was perfused inside the crosslinked print without any leakage. **(e)** Vascular-like networks with branch points can connect channels with varying inner diameters, with both parent (2.5 mm diameter) and daughter (1.5 mm diameter) vessels. Red dye was perfused inside the crosslinked print without any leakage. **(f)** Brightfield microscopy images of alginate, GelMA, and fibrin channels and **(g)** their outer shell thickness as a function of diffusion time. **(h)** FRAP analysis for freely diffusing fluorescent dextrans of varying molecular weights (0.7 - 250 kDa) in alginate, GelMA, and fibrin gel precursor support materials. **(i)** Confocal microscopy images of a fluorescent dextran (FITC-dextran, MW = 10 kDa) diffusing from a single printed filament in a GelMA support material as a function of diffusion time. **(j)** Experimental measurements of diffusion are in close agreement with simulation results of predicted diffusion obtained with finite element analysis. **(k)** GelMA shell thickness is dependent on the concentration of the reaction-initiator (LAP) in the sacrificial ink.

While the inner diameter of GUIDE-3DP channels is controlled through parameters during the printing process (*i*.*e*., applied pressure and speed), the outer diameter of the channel shell is controlled by the diffusion time after printing, during which the reaction-initiators diffuse away from the sacrificial ink into the surrounding gel precursor support material. With increasing diffusion time into the support material, the reaction-initiator is able to reach a farther distance away from the sacrificial ink, thus increasing the thickness of the outer shell (**Figures 3f-g; S3a-c**). Therefore, the relationship between the post-printing diffusion time and outer diameter thickness is dependent on the diffusion rate of the reaction-initiator within the gel precursor support material, which can be modeled as Fickian diffusion. The diffusion rate can be predicted by estimating the diffusivity of the reaction-initiator through the support material. Here we demonstrate examples for three different gel precursor materials and crosslinking approaches: GelMA as a photocrosslinked material (reaction-initiator: LAP), alginate as a small molecule crosslinked material (reaction-initiator: Ca^2+^), and fibrin as an enzymatically crosslinked material (reaction-initiator: thrombin). Fluorescence Recovery After Photobleaching (FRAP) measurements on solutes of different molecular weights (ranging between 0.7 and 250 kDa) were used to estimate the diffusivity for the corresponding reaction-initiators (**Figures 3h; S3d**). For reaction-initiators with lower molecular weights (MW), the higher diffusivity allows the reaction-initiator to diffuse faster (*e*.*g*., Ca^2+^ for alginate crosslinking, MW = 40 Da, D_est_ ∼ 800 μm^2^/s *vs*. thrombin for fibrin crosslinking, MW = 36 kDa, D_est_ ∼ 4 μm^2^/s). This leads to a faster rate of outer diameter increase for high-diffusivity material systems (*i*.*e*. Ca^2+^/alginate) compared to low-diffusivity systems (*i*.*e*. thrombin/fibrin) (**Figure 3g**).

For a given support material/reaction-initiator combination, the Fickian diffusion of the reaction-initiator over time can be monitored and predicted with finite element modeling (FEM). Specifically, we employed FEM to study the diffusion of LAP (MW = 294 Da, D_est_ ∼ 80 μm^2^/s) through a GelMA gel precursor material. By first using fluorescent dextran (FITC-dextran, MW = 1000 Da) as a tracer molecule in the GelMA gel precursor material, we measured the experimental diffusion rate away from the printed sacrificial ink filament (**Figures 3i**). These measurements exhibited excellent agreement with the computational predictions (**Figure 3j**) using the diffusion coefficient obtained from experimental FRAP analysis (**Figure 3h**). Finally, in addition to the diffusion time, one can also control the diffusion rate, and hence the outer shell diameter, by simply changing the concentration of the reaction-initiator in the sacrificial ink. With higher concentrations of LAP encapsulated in the sacrificial ink (tested between 0.25 mM and 1.5 mM LAP), the outer GelMA shell diameter at a given time point increases (**Figure 3k**). With these strategies, both the inner channel diameter and the outer shell diameter of the GUIDE-3DP channels can be independently controlled.

Of the three demonstrated crosslinking approaches for solidifying the channel into a robust gel, photocrosslinking requires a light-initiation step to induce the reaction, effectively decoupling the diffusion and reaction steps and allowing fabrication of more complex structures (**Figure 4**). As a first example, we demonstrate the ability to control the intersection of two GelMA channels (**Figures S4a; 4a**). By printing two LAP-containing sacrificial ink filaments that contact each other within a GelMA support material, a continuously smooth interface is created. Following UV exposure, this results in the fabrication of a branched network with an open, seamless, leak-proof interface (**Figure S4a**). Fabricating these types of free-standing, intersecting channel structures has previously proved difficult with other technologies.^[29,30]^

**Figure 4.**
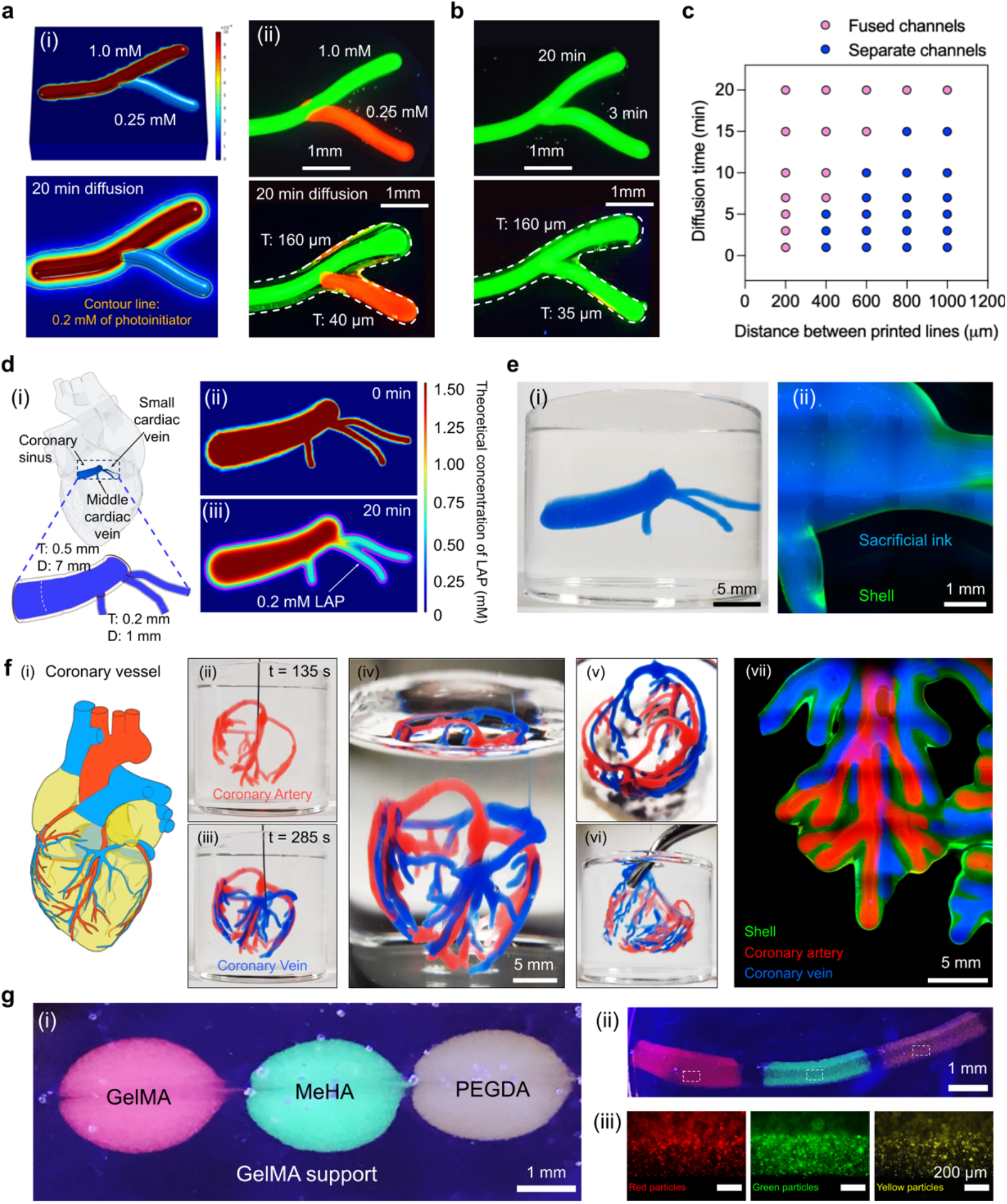
Complex perfusable networks are fabricated with the GUIDE-3DP method. **(a)** (panel i) A computational model of a bifurcated channel predicts LAP diffusion into GelMA for a structure using two sacrificial inks with different LAP concentrations. (panel ii) GUIDE-3DP printing of the model in (i) confirmed fabrication of channels with differing shell thickness connected at an open branch point. **(b)** Fabrication of channels with differing shell thickness is also achieved by controlling the LAP diffusion time from a single sacrificial ink. **(c)** Neighboring, parallel channels can either fuse together to form a single bulk structure with two internal channels or remain as two distinct, separate channels. **(d)** Construction of a print mimicking the coronary sinus connected to small and middle cardiac veins (panel i). The theoretical concentration of LAP initially (panel ii) and 20 mins after 3D printing (panel iii). **(e)** Photo of printed coronary sinus and cardiac veins (panel i) and close-up fluorescent micrograph (panel ii) showing the GelMA shell (green) and sacrificial ink (blue). **(f)** To fabricate a model of the coronary vessels (panel i), the artery and vein networks were printed sequentially (panels ii and iii). Side and top views of the printed coronary vessel structure (panels iv and v). The printed coronary vessel structure can be picked up with forceps (panel vi). Close-up fluorescent micrograph showing the GelMA shell (green) around the arterial network (red) and the venous network (blue) (panel vii). **(g)** Multiple photocrosslinkable materials (GelMA, MeHA, and PEGDA) can be fabricated into a single channel structure using a print-in-print method. Individual regions of labeled GelMA (red), MeHA (green), and PEGDA (yellow) are first printed within a GelMA support material (panel i). All three gel precursors are crosslinked with the same reaction-initiator (LAP) to form a continuous channel (panels ii and iii).

Furthermore, we can design open networks in which different branches have unique, user-specified diameters. As described above, two strategies can be used to control the outer diameter of printed GUIDE-3DP channels: varying reaction-initiator concentration or diffusion time (**Figures 3f-k**). To demonstrate how these strategies can be employed in a more complex structure, we varied the concentration of photoinitiator within different, connected ink filaments. By modeling the photoinitiator diffusion (**Figure 4a, panel i**), the concentrations required to achieve the desired outer channel diameters can be determined. In this way, connected channels of different channel thicknesses (160 μm for the top branch with 1.0 mM LAP; 40 μm for the bottom branch with 0.25 mM LAP) are constructed (**Figure 4a, panel ii**). This type of structure (*i*.*e*., varied channel diameters connected at an open branch point) can also be achieved with a complementary approach in which the diffusion time of the photoinitiator from the ink is varied. In this demonstration, one filament (bottom) is printed 17 min after the first filament (top), such that the photoinitiator in the first filament has had more time to diffuse out, leading to a thicker channel (160 μm for the top branch with 20 min LAP diffusion time; 35 μm for the bottom branch with 3 min LAP diffusion time) (**Figure 4b**). The ability to fabricate these perfusable networks with open branch points and bespoke channel sizes is uniquely enabled by material systems in which the diffusion and reaction processes are decoupled, demonstrated here with GelMA/LAP as an exemplary photocrosslinkable material.

When two neighboring, parallel lines are printed, they can either fuse to form a single bulk structure with two internal channels or remain as two distinct, separate channels (**Figure S4b**). Specifically, the fabrication of cohesive, fused channels is facilitated by shorter distances between the printed lines and longer diffusion times (**Figure 4c**). These predictions allow for the design of multiple perfusable networks fused into a cohesive structure, which may be advantageous when fabricating complex constructs that require manipulation (*e*.*g*. for implantation) due to the increased structural rigidity.

Having established these physical principles of GUIDE-3DP with photoactive crosslinkers (**Figure 4a-c**), we next sought to demonstrate fabrication of physiologically relevant structures. As a first example, we selected a model of the human coronary sinus and cardiac veins, which have differing internal and outer diameters (**Figure 4d, panel i**).^[31,32]^ COMSOL modeling of LAP diffusion through GelMA informed the diffusion time required to fabricate these branched channels of desired geometry (**Figure 4d, panels ii and iii**). The creation of this tubular architecture with open intersections of multiple smaller channels was validated with a GelMA print, which had sufficient structural integrity to be removed from the support material for imaging (**Figure 4e**).

Using the GUIDE-3DP strategy with photocrosslinkable materials also allows us to design two or more distinct perfusable networks within the same structure (**Figures S4c; 4f**). This is enabled by formulating the sacrificial ink to be thixotropic and self-healing, which allows the nozzle to pass through the printed path of the first network without causing permanent deformation while printing the path of the second network. In this way, two intertwined perfusable channels were fabricated, opening the door to printing of more complex vascular-like networks, which often require the extrusion nozzle to pass through ink that has been previously deposited (**Figure S4c**).

As a demonstration, we printed a coronary vascular network geometry within a GelMA support using cardiac structural data from the National Institutes of Health (NIH) 3D Print Exchange (**Figure 4f**). This coronary vessel structure was built by first printing the coronary artery structure (red ink) and then the coronary vein structure (blue ink) (**Figure 4f, panels ii and iii**). Both coronary structures were stable during printing and exhibited no visible changes after UV crosslinking. Crosslinking was performed 20 min after finishing the printing process, demonstrating the short working times achievable with our GUIDE-3DP method (**Figure 4f, panels iv and v**). The coronary vessel-like print demonstrated high structural integrity, allowing it to be removed from the uncrosslinked gel precursor support material with forceps for rinsing with saline (**Figure 4f, panel vi**). Confocal microscopy images show that the UV-crosslinked shell structures stained by FITC-dextran homogeneously surround the printed vessels (**Figure 4f, panel vii**). We observed excellent extrusion consistency and structural stability throughout the 3D printing process, which was evident from the high lateral and axial uniformity observed across the coronary artery and coronary vein structures.

With the GUIDE-3DP strategy, different materials that utilize the same reaction-initiator may also be crosslinked together into a cohesive print. As a demonstration, we fabricated three photocrosslinkable materials into a single, perfusable structure (**Figure 4g**). Pre-labeled GelMA, MeHA, and PEGDA were first 3D printed into distinct regions within a larger, unlabeled GelMA support matrix; then upon printing a single, LAP-containing sacrificial ink, a continuous channel with different material segments was formed.

Finally, the GUIDE-3DP strategy allows for not only fabrication of channels with biologically relevant geometries but also integration with living cells. As the hydrogel materials used in this study for GUIDE-3DP are highly cytocompatible^[33]^, tailored vessel structures can be seeded with human umbilical vein endothelial cells (HUVECs) to achieve an *in vitro* cell-lined blood vessel model. Customizable vessels with tunable dimensions and multiple furcation points are highly relevant as models for stenosis or vessel calcification.^[34,35]^

Successful seeding of HUVECs was demonstrated within GelMA channels fabricated with GUIDE-3DP (**Figure 5**). The HUVECs were introduced into the hollow channels with perfusion, and adherent cells showed high cell viability 3 days after seeding (**Figure 5a-b**). As a demonstration of a more complex vasculature-like structure, we fabricated a branched channel network (**Figure 5c**). The channel, which consisted of a thicker parent vessel (inner diameter: 2 mm) with four smaller daughter vessels (inner diameters: 1.0-1.5 mm), showed a continuously open lumen that was fully perfusable without leakage (**Figure 5c, panel i**). Therefore, the lining of the vessel lumen with viable HUVECs was possible (**Figure 5c, panels ii, iii, iv**).

**Figure 5.**
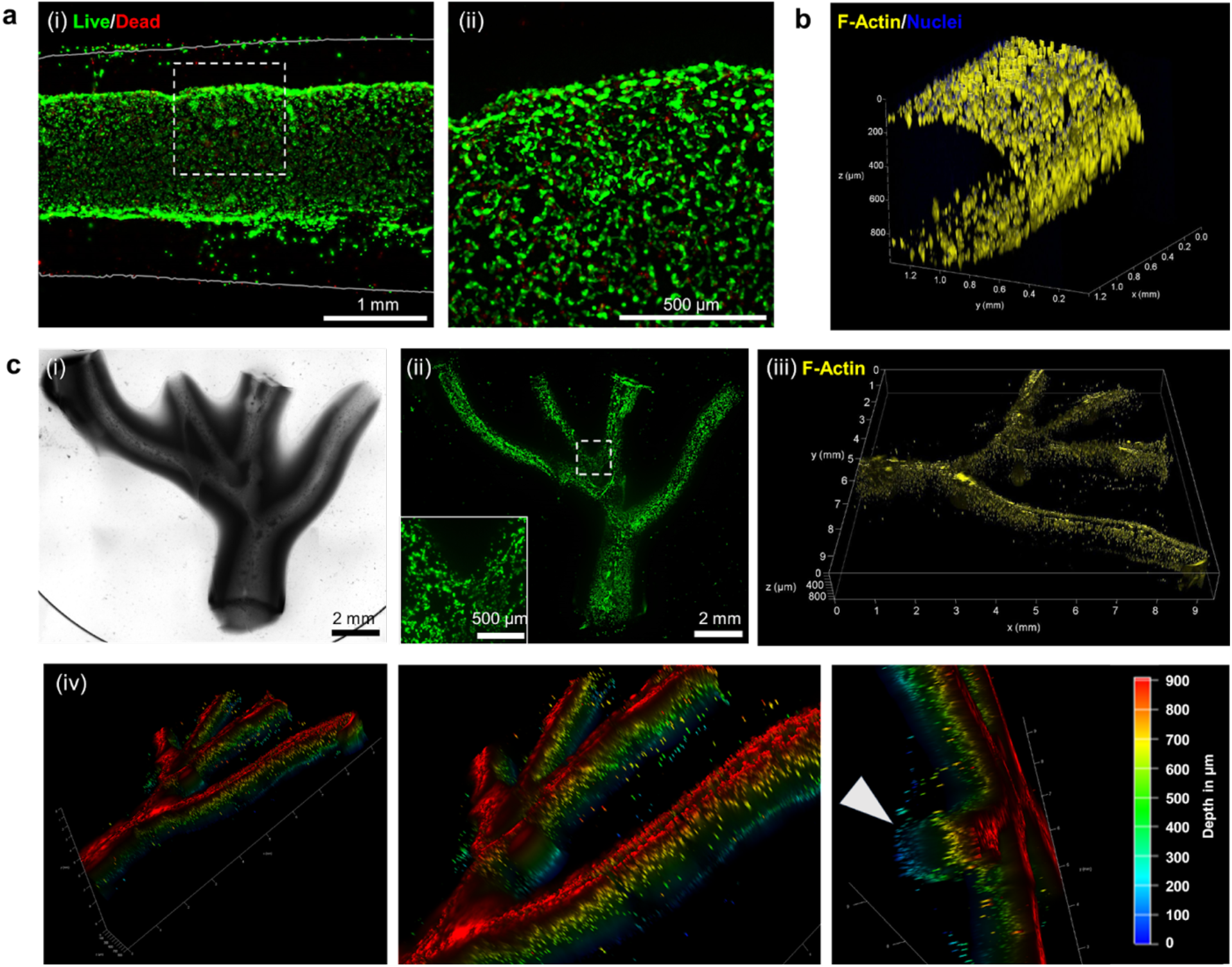
The inner lumens of perfusable vessels printed with GUIDE-3DP can be seeded with endothelial cells. **(a)** Representative fluorescence images of a GelMA channel 3 days post-seeding: (i) image of viable (green, calcein AM) and dead (red, ethidium homodimer-1) HUVECs within the inner lumen (diameter = 2 mm), with the outer vessel wall (thickness = 1 mm) indicated by the solid gray lines; (ii) magnified view of cells at the vessel wall interface, with the acquisition position indicated by the dashed white box in (i). **(b)** 3D reconstruction of the HUVEC-lined vessel stained for F-actin cytoskeleton (yellow) and nuclei (blue). **(c)** Representative images of a HUVEC-lined, furcated GelMA structure: (i) brightfield image; (ii) fluorescence image of viable HUVECs (green, calcein AM) and (inset) magnified view at a channel furcation point; (iii) 3D reconstruction of the furcated channel network stained for F-actin (yellow); and (iv) depth-coded images with white arrow indicating the patent, circular vessel lumen.

As endothelial cells are known to adhere well on fibronectin matrices *in vitro*^[36]^, we incorporated 0.05 wt% fibronectin within the GelMA gel precursor support material prior to printing to demonstrate versatility in the biochemistry of GUIDE-3DP structures. While the viability of HUVECs was similar (around 70 % live cells) for seeding within GelMA-only or GelMA-fibronectin prints (**Figure S5**), the HUVECs exhibited greater spreading and expression of the endothelial adhesion molecule VE-cadherin along the inner lumens of GelMA-fibronectin channels (**Figure S6**). The enhanced cellular spreading due to the added fibronectin is demonstrated by the increased overall cytoskeletal area per cell (261 ± 127 µm^2^ for GelMA-fibronectin *vs*. 135 ± 84 µm^2^ for GelMA-only). Thus, the GUIDE-3DP strategy allows for the integration of complex, branching, perfusable channels with living cells and is amenable to the addition of biomolecules in the gel precursor support material for future optimization of cell function.

## 3. CONCLUSIONS

We report a new 3D printing strategy termed Gelation of Uniform Interfacial Diffusant in Embedded 3D Printing (GUIDE-3DP) for fabricating bespoke, perfusable networks of self-supporting, interconnected channels. To achieve user-specified channel dimensions, this technique leverages the predictable diffusion of reaction-initiators released from sacrificial inks printed within the hydrogel precursor. The inner and outer diameters of vessels can be independently varied within a single print: inner diameters are controlled by the printing pressure and speed, while outer diameters are controlled by the reaction-initiator concentration and diffusion time. Finite element modeling can be used to relate the desired dimensions of a channel network to the printing parameters. Importantly, multiple channels can be seamlessly connected at branch points, and multiple networks of channels can be integrated into a single, cohesive, self-supporting print. We created a library of seven hydrogel precursor materials (gelled *via* photocrosslinking, small molecule crosslinking, or enzymatic crosslinking) and their corresponding reaction-initiators to highlight the versatility of this technology. Finally, we demonstrated the integration of the GUIDE-3DP approach with living endothelial cells that lined the inner lumens of blood vessel-like channels. Altogether, this technology represents an important advance toward the fabrication of self-supporting, physiologically relevant networks with intricate and perfusable geometries.

## 4. EXPERIMENTAL SECTION

### Gel precursor support materials preparation

For the gelatin methacryloyl (GelMA) gel precursor material, GelMA was synthesized using a protocol similar to those described previously.^[37]^ Gelatin from cold water fish skin (Sigma-Aldrich) was dissolved at 20 wt% in 0.1 M carbonate-bicarbonate (CB) buffer at 37 °C overnight. The pH was adjusted to 10. 83 µL of methacrylic anhydride (MAA, 94%, Sigma-Aldrich) per gram of gelatin was added dropwise while stirring at 100-200 rpm. The reaction was allowed to proceed at 70 °C for 2 h. GelMA was precipitated and collected after pouring the reaction mixture into ∼3x reaction volume of ethanol. After drying the GelMA (> 24 h), it was dissolved in deionized water at ∼80 °C for at least 1 h while stirring to remove residual ethanol. The GelMA concentration was determined as percentage of dry mass. To prepare the support material, respective stock solutions of GelMA and Aristoflex Aristoflex Ammonium Acryloyldimethyltaurate/VC Copolymer (AVC; Lotioncrafter) were blended to achieve the desired final concentrations.

For the poly(ethylene glycol) diacrylate (PEGDA) support material, 10 wt% of poly(ethylene glycol) diacrylate (PEGDA, MW: 20 kDa) was dissolved in PBS. The 10 wt% PEGDA solution was mixed in a 1:1 ratio with a 4 wt% AVC stock solution to form the 5 wt% PEGDA + 2 wt% AVC support material.

For the methacrylated hyaluronic acid (MeHA) support material, sodium hyaluronic acid (HA, Lifecore Biomedical, 40kDa Mw) was dissolved in deionized water at 1 wt%. Methacrylic anhydride (MAA, 94%, Sigma-Aldrich) was slowly added to the solution. The pH was adjusted to 8–9 with 6 N sodium hydroxide solution and the solution was gently stirred at 4 °C for 18 h, protected from light. The MeHA was subsequently precipitated by ethanol (95%) and dialyzed against deionized water using a cellulose acetate tube (molecular weight cut-off: 12 kDa) for three days, followed by lyophilization. The 2 wt% MeHA solution was then mixed in a 1:1 ratio with a 4 wt% AVC stock solution to form the 1 wt% MeHA + 2 wt% AVC support material.

For the polyacrylamide (PAAm) support material, 40 wt% of acrylamide (AAm, Sigma-Aldrich) and 0.2 wt% of N,N′-methylenebis(acrylamide) (Sigma-Aldrich) was dissolved in deionized water. They were mixed with a 4 wt% AVC stock solution to form the 20 wt% acrylamide + 0.1 wt% N,N′-methylenebis(acrylamide) + 2 wt% AVC support material.

For the alginate support material, alginic acid sodium salt from brown algae (Sigma-Aldrich) was dissolved at a concentration of 4 wt% in PBS. The 4 wt% alginate solution was then mixed in a 1:1 ratio with a 4 wt% AVC stock solution to form the 2 wt% alginate + 2 wt% AVC support material.

For the fibrin support material, bovine fibrinogen (MP Biomedicals) was dissolved at a concentration of 10 wt% in PBS, and then mixed with a 4 wt% AVC stock solution to form the 5 wt% fibrinogen + 2 wt% AVC support material.

For the gelatin support material, gelatin from cold water fish skin (Sigma-Aldrich) was dissolved at 20 wt% in PBS, and then mixed with a 4 wt% AVC stock solution to form the 10 wt% gelatin + 2 wt% AVC support material.

### Sacrificial ink preparation

To prepare the gelatin microparticle ink used in acellular printing experiments, gelatin microparticles were synthesized by complex coacervation, as previously described.^[37]^ Briefly, a solution of 6.4 wt% gelatin type A (300 bloom, Sigma-Aldrich), 0.5 wt% Pluronic F-127 (Sigma-Aldrich), and 0.2 wt% gum arabic (Alfa Aesar) in DI water was heated in a microwave until bubbling. Ethanol was added at 1 mL per gram of gelatin solution while stirring at 500 rpm. The mixture was then stirred overnight at 500 rpm at room temperature. The gelatin microparticles were collected and washed three times with DI water using a Büchner funnel, and the resultant slurry was compacted *via* vacuum filtration. The concentration of microparticles in the jammed slurry was determined as the percent dry mass after drying aliquots in a vacuum oven for 2 h at 60 °C. Prior to printing, the microparticle concentration was adjusted to 4-10 wt% using a solution of crosslinking initiator in PBS (or HBSS for an alginate support material) such that the desired crosslinking initiator concentration was achieved (**Table S2**). The reaction-initiators included in the inks were calcium chloride dihydrate (Millipore) for an alginate support material, ammonium persulfate (APS, Sigma-Aldrich) for a PAAm support material, bovine thrombin (MP Biomedicals) for a fibrin support material, and lithium phenyl-2,4,6-trimethylbenzoylphosphinate (LAP, Sigma-Aldrich) for GelMA, MeHA, and PEGDA support materials. For cellular printing experiments, sterile lyophilized gelatin microparticles (LifeSupport, FluidForm Inc.) were rehydrated at a concentration of 10 wt% in sterile, cold PBS containing 5 mM LAP. For the gelatin support material, a sacrificial ink of 24 wt% Pluronic F-127 containing 1 wt% transglutaminase was used instead of gelatin microparticles.

### Rheological characterization

To measure the rheological properties of the gel precursor support materials and sacrificial ink, samples were loaded onto an ARG2 stress-controlled rheometer (TA Instruments) equipped with a 40 mm diameter parallel plate geometry at a 1.0 mm gap height. Samples were subjected to shear stress-sweep experiments over a stress range of 0.1 to 1000 Pa at a frequency of 1 Hz at 25 °C. The viscosity was measured at 25 °C using a range of shear rates from 0.1 to 1000 s^−1^. The step-stress was measured at both a high-magnitude stress (300 Pa) and a low-magnitude stress (0.1 Pa). To characterize the temperature dependence of the gelatin microgel ink properties, temperature sweep experiments were performed from 25 to 37 °C.

### Fluorescence recovery after photobleaching (FRAP)

Alginate, GelMA, and fibrin support materials were prepared with FITC-labeled diffusants (either 1 µg/mL diazide-PEG-FITC, 10 µg/mL of 10, 20, 40, 60, 150, 250 kDa FITC-dextran, Sigma). 150 µL of the diffusant-laden support materials were loaded into a clear bottom, 96-well plate and centrifuged to remove bubbles. FRAP experiments were performed using a confocal microscope (Leica SPE) with 30 s of photobleaching (100 µm x 100 µm area, 488 nm laser, 100% intensity) and 90 s of capture time. Diffusion coefficients for each condition were calculated using the open source MATLAB code “frap_analysis” based on the Hankel transform method.^[38]^

### 3D printing

3D printing was performed with a custom-built dual-extruder bioprinter modified from a MakerGear M2 Rev E plastic 3D printer^[39–41]^ and a custom-built pneumatic printer modified from a Creality Ender 3 3D printer^[42,43]^. Prior to printing, the sacrificial ink was loaded into a 2.5 mL Hamilton Gastight syringe. Support materials were added to well plates or custom-made polycarbonate containers and centrifuged at 4000 rpm for 5 min at room temperature to remove bubbles. Print paths were created using the commercially available Rhinoceros software (Rhinoceros 5.0, Robert McNeel & Associates, Seattle, WA, USA). Using the Rhinoceros software, designed printing paths (2D lines) were segmented with ‘divide’ function to create 1D dots to determine the cartesian coordinates. The cartesian coordinates were saved as a .txt file and translated into G-code to set extrusion rates using CAMotics software. After printing, diffusion of the reaction-initiator was allowed to occur for a specified amount of time (**Table S1**). For photocrosslinkable materials (GelMA, MeHA, and PEGDA), the printed structure was crosslinked at 365 nm for 5 min using a UV lamp. After crosslinking, the structure was removed from the uncrosslinked support material using a metal spatula and washed by gently shaking in a 50 mL conical tube containing PBS (or HBSS for an alginate support materials). Once washed, the structures were placed into fresh PBS (or HBSS), and the sacrificial ink was melted by incubating at 37 °C for gelatin microparticles or at 4 °C for Pluronic F-127. The ends of the structures were cut, either before or after sacrificial ink melting, to allow fluid perfusion.

As preparation for endothelialization, materials and channels were processed in an aseptic environment. The support materials were autoclaved prior to use. To create linear channels (length = 2 cm), a 10 wt% gelatin microparticle ink containing 5 mM LAP was printed into a 20 wt% GelMA + 2.5 wt% AVC support material with and without 0.05 wt% fibronectin. To create the branched structure (total size *x* = 13.7 mm, *y* = 11.7 mm), the same sacrificial ink was printed into a 20 wt% GelMA + 2.5 wt% AVC support material. After printing, the ends of the channels were cut using a sterile scalpel to enable subsequent perfusion seeding. Vessels were incubated in sterile PBS for at least 24 h at 37 °C prior to endothelialization.

### Cell culture and endothelialization of printed structures

Human umbilical vein endothelial cells (HUVEC, PromoCell) were expanded to passage 6 or 7 in endothelial growth media (EGM-2 BulletKit, Lonza), and culture medium was changed every other day. For cell seeding on the inner lumen of 3D printed channels, HUVECs were enzymatically detached in 0.025% Trypsin-EDTA (Gibco) for 5 min at 37 °C and resuspended in EGM-2 media at ∼1 × 10^7^ cells/mL. The cell suspension was then injected via pipette into the vessel lumen (approx. 20 µL ≙ 2 ×10^5^ cells per cm of vessel length). The channels were incubated at 37°C for a total of 80 min, during which two incubation steps were performed on each side, flipping by 180° and injecting 20 µL of fresh cell suspension between each step, to allow HUVEC adhesion to the entire inner lumen. Vessels were incubated in EGM-2 medium after cell seeding, and culture medium was changed every other day.

### Fluorescent staining and imaging

Cell viability in printed structures was assessed after 3 days of culture using Live/Dead staining (Thermo Fisher Scientific). Live cells were stained with calcein AM (2 µM) and dead cells were stained with ethidium homodimer-1 (EthD-1; 4 µM) in PBS for 30 min at 37 °C. The printed structures were then imaged using a Leica THUNDER fluorescence microscope using 2.5X and 10X objectives. Z-stack images (total thickness 800-1000 µm) were processed in Fiji (version 1.53).^[44]^ Cell viability was calculated as the ratio (in %) of viable cells (calcein AM-stained cells) to all cells (sum of calcein AM- and EthD-1-stained cells) with N = 3 replicate printed structures, n = 1211-4133 cells per printed structure.

For immunocytochemistry to observe cell distribution and morphology, samples were fixed for 30 min using 4% paraformaldehyde in PBS, then washed three times with PBS for 15 min each. Cells were permeabilized with 0.1% Triton X-100 in PBS (PBST) for 1 h. Non-specific binding was prevented by blocking in 0.05% Triton X-100, 5% normal goat serum, and 0.5 wt% bovine serum albumin (BSA, Roche) in PBS for 3 hours. Primary antibody anti-VE-cadherin (Rabbit mAb, Cell Signaling Technology, 1:250) diluted in 0.05% Triton X-100, 2.5% normal goat serum, and 0.25 wt% BSA was applied for 1 h at room temperature, followed by four washing steps in PBS for 20 min each. A fluorescently tagged secondary antibody (Goat anti-rabbit, Alexa Fluor 647, Thermo Fisher Scientific) diluted in the same antibody solution was applied for 1 h, followed by three PBS washes for 20 min. Cell nuclei and actin cytoskeleton were stained by incubation with 4’,6-diamidino-2-phenylindole, dihydrochloride (DAPI, Thermo Fisher Scientific, 1 µg/mL) and phalloidin–tetramethylrhodamine B isothiocyanate (TRITC-phalloidin, Sigma-Aldrich, 0.2 µg/mL) in PBST for 1 h at room temperature, followed by three PBS washing steps for 15 min. Samples were imaged using a Leica THUNDER fluorescence microscope. Representative images were processed in Fiji, and cell size was quantified using CellProfiler (version 4.2.5).

### Computational modeling

COMSOL Multiphysics (version 5.6) was employed to simulate the diffusion of the photoinitiator LAP from the printed sacrificial ink filament over time. A three-dimensional finite element model (FEM) was created using a Time Dependent study in the Transport of Diluted Species interface. The model geometry was imported with an STL file designed according to the G-code path, and the diameter was set according to that of the printed filament. To simulate LAP transport, the diffusion coefficient of LAP obtained using FRAP (D_est_ ∼ 80 μm^2^/s) was entered into the model, and the initial concentration of LAP in the sacrificial ink was set according to experimental values. A tetrahedral physics-controlled mesh with the predefined Finer element size was used in all simulations. The concentration profiles of LAP as a function of distance from the ink filament were then calculated for diffusion times ranging between 0 and 30 min.

### Statistical analysis

Results were plotted in GraphPad Prism (version 9.3). For the comparison of cell viability and area, statistical significance was assessed using a two-tailed Mann-Whitney test (ns = not significant; **** p<0.0001). Data are presented as the mean ± the standard deviation (SD) unless specified otherwise.

## ACKNOWLEDGMENTS

S.C.H. acknowledges support from the National Science Foundation (DMR-2103812, CBET 2033302) and the National Institutes of Health (R01-EB027171). L.G.B. acknowledges support from the National Science Foundation Graduate Research Fellowship Program (DGE-1656518). B.C. acknowledges support from the Stanford Knight-Hennessy Scholars program. J.G.R. acknowledges support from the National Science Foundation Graduate Research Fellowship Program (DGE-1656518) and the Stanford Smith Family Graduate Fellowship.

## CONFLICT OF INTEREST

S.S., J.G.R., A.J.S., and S.C.H. are inventors on a provisional patent application (no. 63/338,368) related to this work, submitted by the Board of Trustees of Stanford University.

