## Supplementary Information for "Gelation of Uniform Interfacial Diffusant in Embedded 3D Printing"

**Table S1.** Printing parameters for all constructs presented in this work.

| Figure | Print Description | Gel material* | Diffusion time (min) | Printing speed (mm/s) | Extrusion rate (µl/s) or printing pressure (psi) |
|---|---|---|---|---|---|
| 1b | Single linear line | GelMA (10%) | 5 min | 5 mm/s | 15 µl/s |
| 1c (i) | Single linear line | GelMA (10%) | 20 min | 5 mm/s | 4 µl/s |
| 1c (ii) | Single linear line | GelMA (10%) | 1 min | 5 mm/s | 4 µl/s |
| 1c (iii) | Fused 8 lines | GelMA (10%) | N/A | 5 mm/s | 4 µl/s |
| 1c (iv) | Fused spiral line | GelMA (10%) | N/A | 5 mm/s | 4 µl/s |
| 1d | Winding lines | GelMA (10%) | 10 min | 5 mm/s | 4 µl/s |
| | Winding line | MeHA (1%) | 10 min | 5 mm/s | 4 µl/s |
| | Winding line | PEGDA (5%) | 10 min | 5 mm/s | 4 µl/s |
| | Winding line | Alginate (2%) | 5 min | 5 mm/s | 4 µl/s |
| | Winding line | PAAm (20%) | 5 min | 5 mm/s | 4 µl/s |
| | Winding line | Fibrin (5%) | 20 min | 5 mm/s | 4 µl/s |
| | Winding line | Gelatin (10%) | 20 min | 5 mm/s | 4 µl/s |
| 2g (i) | Letters ("GelMA") | GelMA (10%) | N/A | 5 mm/s | 0.4 µl/s |
| 2g (ii) | Letters ("MeHA") | MeHA (1%) | N/A | 5 mm/s | 0.4 µl/s |
| 2g (iii) | Letters ("PAam") | PAAm (20%) | N/A | 5 mm/s | 0.4 µl/s |
| 2g (iv) | Letters ("PEGDA") | PEGDA (5%) | N/A | 5 mm/s | 0.4 µl/s |
| S1a, S2a | Single linear line, right angle | GelMA (10%) | N/A | 5 mm/s | 4 µl/s |
| 3a | Single linear line | GelMA (10%) | N/A | 5 mm/s | Variable pressure |
| 3b | Lines | GelMA (10%) | 5 min | 5 mm/s | Variable pressure |
| 3c | Lines | GelMA (10%) | 5 min | Variable speed | 25 psi |
| 3d | Intestinal | GelMA (10%) | 5 min | 2 mm/s and 1 mm/s | 40 psi |
| 3e | Vasculature | GelMA (10%) | 5 min | 3 mm/s and 4 mm/s | 40 psi |
| 3f, S3 | Single linear line | Alginate (2%) | Variable times | 5 mm/s | 4 µl/s |
| | Single linear line | GelMA (10%) | Variable times | 5 mm/s | 4 µl/s |
| | Single linear line | Fibrin (5%) | Variable times | 5 mm/s | 4 µl/s |
| 3i, 3j | Single linear line | GelMA (10%) | Variable times | 5 mm/s | 4 µl/s |
| 4a, 4b | Branched lines | GelMA (10%) | 20 min | 5 mm/s | 1 µl/s |
| 4e | Coronary sinus | GelMA (10%) | 20 min | 5 mm/s (10 times) | 4 µl/s |
| | Small cardiac vein | GelMA (10%) | 3 min | 5 mm/s (1 time) | 4 µl/s |
| 4f | Coronary vessel network | GelMA (10%) | 20 min | 5 mm/s | 4 µl/s |
| 4g | Single linear line | GelMA (10%), MeHA (1%), PEGDA (5%) | 5 min | 5 mm/s | 4 µl/s |
| S4a | Intersection | GelMA (10%) | 5 min | 5 mm/s | 4 µl/s |
| S4b | Fused channels | GelMA (10%) | 5 min | 5 mm/s | 4 µl/s |
| S4c | Axial vessel and helix | GelMA (10%) | 5 min | 5 mm/s | 4 µl/s |
| 5a | Single linear line | GelMA (20%) | 5 min | 5 mm/s | 8 µl/s |
| 5b | Branched vascular-like channel | GelMA (20%) | 5 min | 5 mm/s | 8 µl/s |
| S5a, b | Single linear line | GelMA (20%), AVC (2.5%) fibronectin (0.05%) | 5 min | 5 mm/s | 8 µl/s |
| S6a, b | Single linear line | GelMA (20%), AVC (2.5%), fibronectin (0.05%) | 5 min | 5 mm/s | 8 µl/s |

*All gel materials included 2 wt% Aristoflex AVC unless specified otherwise. 3D printing was performed with a 27G nozzle in all cases.

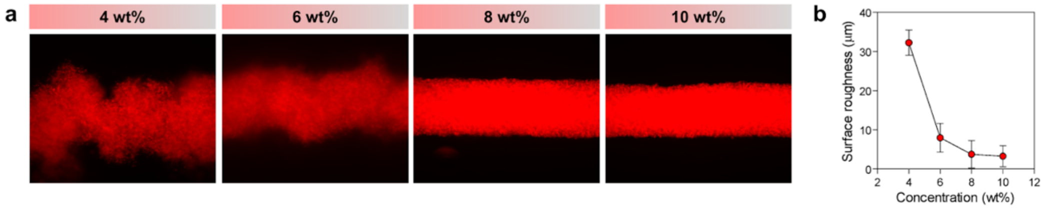


**Figure S1.** Surface roughness of 3D printed gelatin microparticle ink filaments of varying concentrations. **(a)** The concentration of gelatin microparticles (4-10 wt%) affects the fidelity of printed lines within a GelMA gel precursor support material (10 wt% GelMA, 2 wt% AVC). **(b)** The surface roughness of the printed gelatin microparticle ink filament decreases with increasing concentration of the gelatin microparticles, from 4-10 wt%.

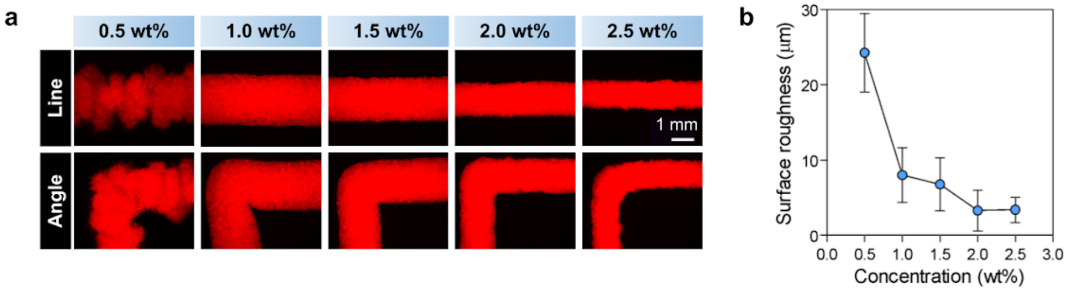


**Figure S2.** Surface roughness of 3D printed gelatin microparticle ink filaments within a 10 wt% GelMA gel precursor support material with varying concentrations of Aristoflex AVC. **(a)** The concentration of the viscosity modifier Aristoflex AVC (0.5-2.5 wt%) added to 10 wt% GelMA affects the fidelity of printed lines and angles (8 wt% gelatin microparticles). **(b)** The surface roughness of the printed gelatin microparticle ink filament (8 wt%) decreases with increasing concentration of AVC within the GelMA, from 0.5-2.5 wt%.

**Table S2.** Formulations of the gel precursor support materials and sacrificial inks used for GUIDE-3DP.

| Mechanism | Support material | | | Sacrificial ink | | |
|---|---|---|---|---|---|---|
| | Hydrogel | Co-initiator | Viscosity modifier | Gelatin microgels | Pluronic F-127 | Reaction-initiator |
| Small molecule crosslinking | Alginate (2 wt%) | - | Aristoflex AVC (2 wt%) | 8 wt% | - | $CaCl_2$ (1 wt%) |
| | PAAm (20 wt%) | TEMED (20 mM) Bis-acrylamide (0.1 wt%) | Aristoflex AVC (2 wt%) | 8 wt% | - | APS (20 mM) |
| Photocrosslinking | GelMA (10 wt%) | - | Aristoflex AVC (2 wt%) | 8 wt% | - | LAP (2 mM) |
| | MeHA (2 wt%) | - | Aristoflex AVC (2 wt%) | 8 wt% | - | LAP (2 mM) |
| | PEGDA (5 wt%) | - | Aristoflex AVC (2 wt%) | 8 wt% | - | LAP (2 mM) |
| Enzymatic crosslinking | Fibrin (5 wt%) | - | Aristoflex AVC (2 wt%) | 8 wt% | - | Thrombin (500 $Uml^{-1}$) |
| | Gelatin (10 wt%) | - | Aristoflex AVC (2 wt%) | - | 24 wt% | Transglutaminase (1 wt%) |

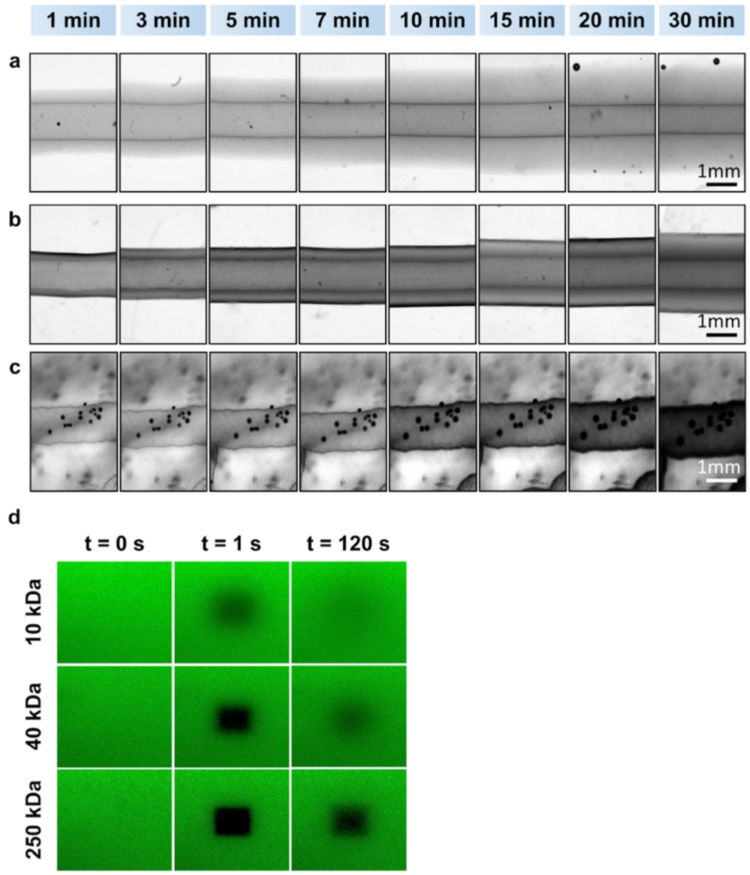


**Figure S3.** The outer diameters of GUIDE-3DP channels increase with greater reaction-initiator diffusion times within **(a)** alginate, **(b)** GelMA, and **(c)** fibrin gel precursor support materials. **(d)** Representative FRAP images for FITC-dextrans (MW: 10 kDa, 40 kDa, and 250 kDa) in a GelMA support material, showing pre- and post-bleaching frames.

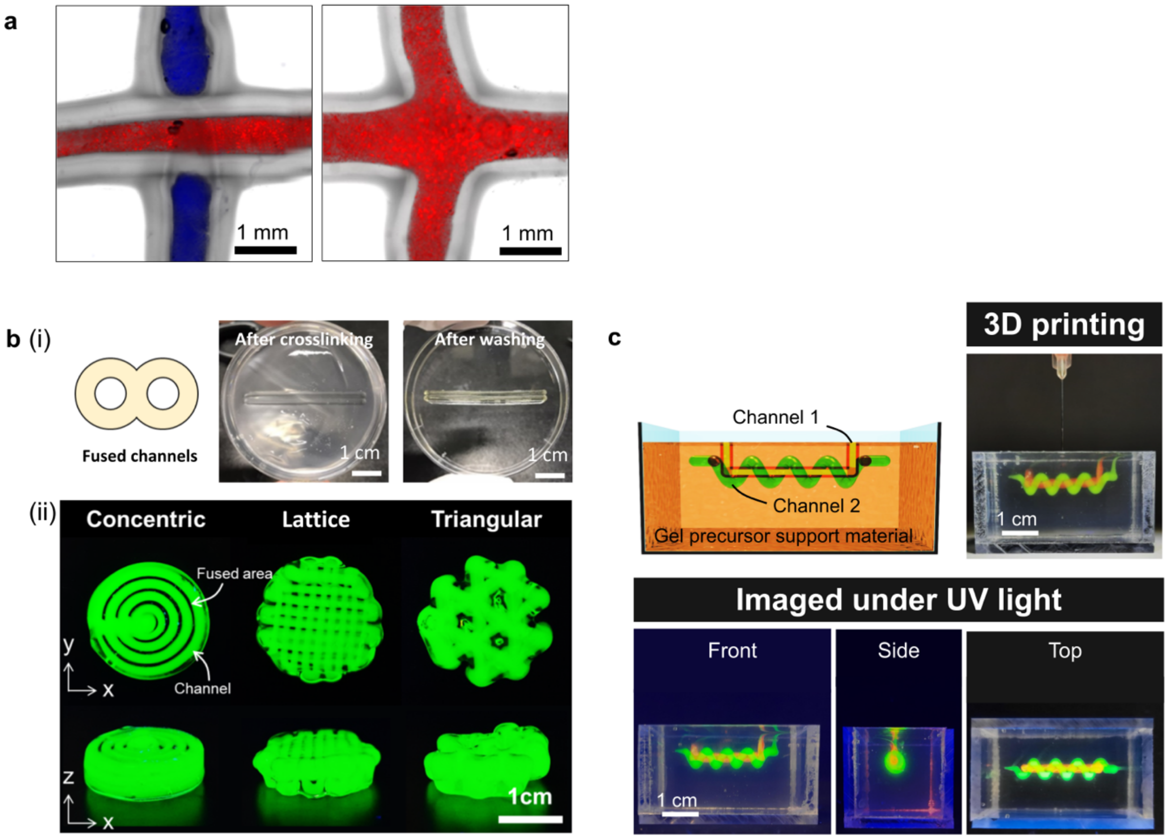


**Figure S4. (a)** GUIDE-3DP allows for open branch points at the intersection of two printed channels for photocrosslinkable materials, since the photocrosslinking process can be initiated on-demand upon exposure to UV light. When the GelMA shell is crosslinked immediately after each single line is printed, distinct, non-intersecting channels are formed (left panel). However, if photocrosslinking is initiated only after printing of both lines is completed, an open branch point is formed (right panel). **(b)** Neighboring, parallel channels can fuse together to form a single bulk structure with two internal channels (panel i). Various fused channel networks can be printed, including concentric, lattice, and triangular designs (panel ii). **(c)** Multiple, intertwined perfusable channels can be integrated within a single GelMA print, demonstrated here by a straight channel 1 (orange) wrapped by a spiral channel 2 (green). The sacrificial inks for both channels were printed prior to photocrosslinking.

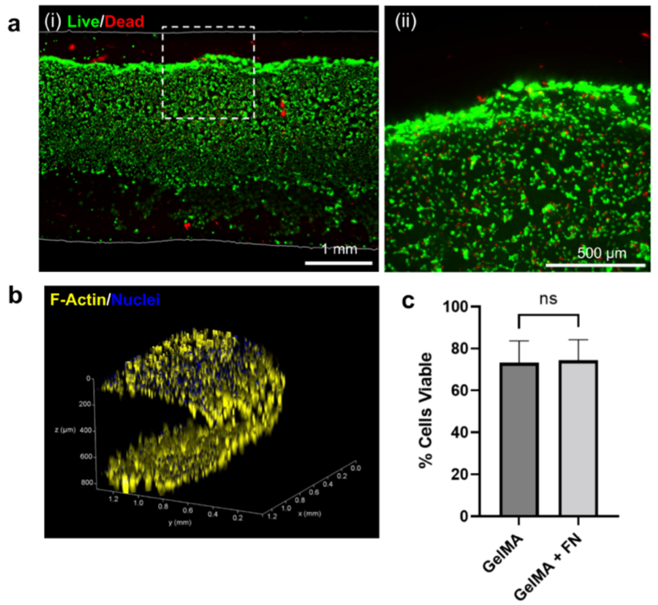


**Figure S5.** Endothelial cells seeded within GelMA-fibronectin prints. **(a)** Representative fluorescence images of a GelMA-fibronectin channel 3 days after seeding: (i) fluorescence image of viable (green, calcein AM) and dead (red, ethidium homodimer-1) HUVECs within the inner lumen (diameter = 1.4 mm), with the outer vessel wall (thickness = 0.8 mm) indicated by solid gray lines; (ii) magnified view of cells at the vessel wall interface, with the acquisition position indicated by dashed white box in (i). **(b)** 3D reconstruction of the HUVEC-lined vessel stained for F-actin cytoskeleton (yellow) and nuclei (blue). (**c**) HUVEC viability on GelMA and GelMA + fibronectin (FN) vessels 3 days post-seeding, ns: not significant.

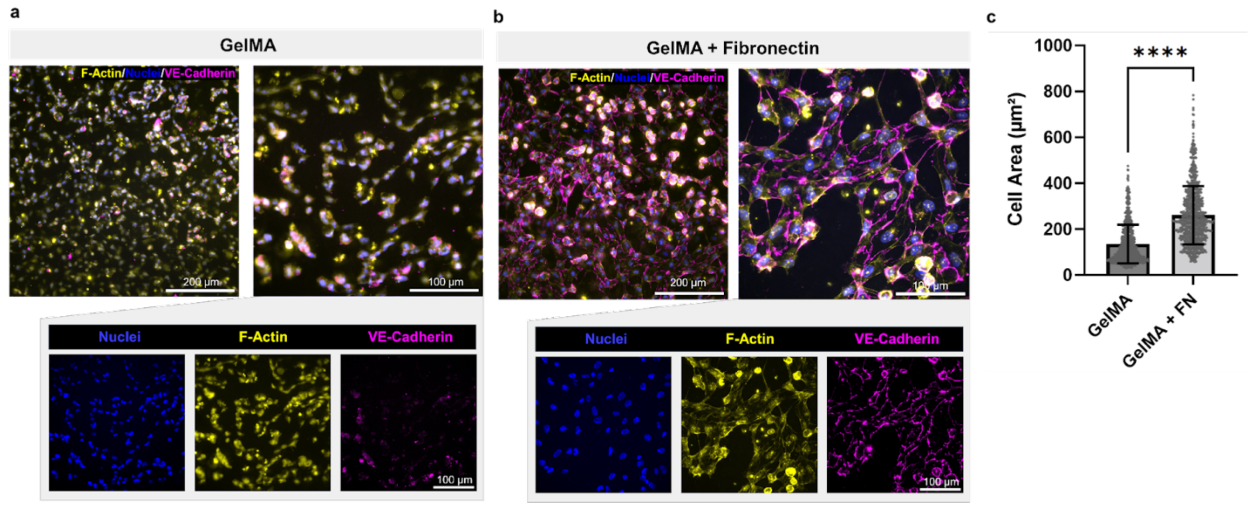


**Figure S6.** Immunocytochemistry of endothelial cells seeded within GUIDE-3DP prints. Representative images of HUVEC-lined (**a**) GelMA and (**b**) GelMA + fibronectin vessels stained for nuclei (blue), F-actin (yellow), and VE-cadherin (magenta) 3 days post-seeding. (**c**) HUVEC cell areas on GelMA and GelMA + fibronectin (FN) vessels 3 days post-seeding, ****$p$<0.0001.
